# ScaleSC: A superfast and scalable single cell RNA-seq data analysis pipeline powered by GPU

**DOI:** 10.1101/2025.01.28.635256

**Authors:** Wenxing Hu, Haotian Zhang, Yu H. Sun, Shaolong Cao, Jake Gagnon, Zhengyu Ouyang, Yuka Moroishi, Baohong Zhang

**Affiliations:** Research Department, Biogen, Inc., 225 Binney St, Cambridge, 02142, MA, USA; Data Science, BioInfoRx, Inc., Madison, 53719, WI, USA

**Keywords:** single cell, scRNA-seq, Scanpy, Seurat

## Abstract

The rise of large-scale single-cell RNA-seq data has introduced challenges in data processing due to its slow speed. Leveraging advancements in GPU computing ecosystems, such as CuPy, and building on Scanpy and rapids-singlecell package, we developed ScaleSC, a GPU-accelerated solution for single-cell data processing. ScaleSC delivers over a 20x speedup through GPU computing and significantly improves scalability, handling datasets of 10–40 million cells with over 1000 batches by overcoming the memory bottleneck on a single A100 card- far surpassing rapids-singlecell’s capacity of processing only 1 million cells without multi-GPU support. We also resolved discrepancies between GPU and CPU algorithm implementations to ensure consistency. In addition to core optimizations, we developed new advanced tools for marker gene identification, cluster merging, and more, with GPU-optimized implementations seamlessly integrated. Designed for ease of use, the ScaleSC package is compatible with Scanpy workflows, requiring minimal adaptation from users. The ScaleSC package (https://github.com/interactivereport/ScaleSC) promises significant benefits for the single-cell RNA-seq computational community.

## Introduction

Recent advances of single cell RNA sequencing (scRNA-seq) technology enable researchers to study the gene expression profiles of individual cells. Unlike bulk RNA sequencing, which averages gene expression across a population of cells, scRNA-seq captures the heterogeneity within a cell population, offering unprecedented insights into cell types, states, and dynamic biological processes.

Spatial transcriptomics (ST) is another innovative technology that combines gene expression profiling with spatial information, allowing researchers to understand how cellular interactions and tissue organization influence gene expression. Emerging techniques integrate single-cell resolution with spatial precision, such as 10x Genomics Visium HD and Xenium, enabling researchers to resolve cellular-level insights within the tissue landscape. These approaches provide a multidimensional view of tissues, bridging molecular and structural biology.

Both scRNA-seq and ST operate at single-cell resolution, producing exceptionally large scale datasets. ScRNA-seq datasets are highly dimensional, encompassing both numerous observations, such as millions of cells, and a vast array of features, such as 30,000 genes. Analyzing scRNA-seq data is a complex, multi-step process aimed at extracting meaningful biological insights from the transcriptomes of individual cells while removing unwanted noise and batch effects. The data analytical workflow includes quality control (QC), normalization, dimensionality reduction, batch effect correction, clustering, cell type annotation, and various downstream analyses. These steps help mitigate unwanted noise and extract biologically relevant signals from cleaned cells and cell types.

The generated scRNA-seq data, typically delivered as a cell-by-gene matrix, is massive due to the high dimensionality of 20-50k genes and millions of cells. This makes the analytical steps computationally intensive in terms of both space (memory) and time cost. As a result, analyzing large scale scRNA-seq datasets requires high-performance servers with a huge memory capacity and is time-consuming. To address these computational challenges, specialized tools have been developed to manage the scalability of the data and reduce computational costs. These tools focus on reducing memory usage, parallelizing workflows, and optimizing algorithms for efficient, high-throughput analysis. For example, HDF5 https://www.hdfgroup.org/solutions/hdf5/, a dedicated package for fast and flexible I/O for large and complex data [1], is used to support hierarchical metadata structures and out-of-memory operations. Anndata [2] is a dedicated tool designed to effectively manage large cell*×*gene matrices. Both ‘in memory’ and ‘on disk’ data loading are supported in Anndata. SciPy [3] supports sparse matrices to further reduce memory storage and speed up computing operations, by leveraging the high sparsity of cell*×*gene matrices. Highly variable genes (HVG) [4, 5] are used to select features for dimension reduction, reducing from 20-50k features to 1-5k, by excluding genes with small variations. Principal component analysis (PCA) is used subsequently to further reduce feature dimension from 1-5k to 20-200.

Scanpy [6], https://Scanpy.readthedocs.io/en/stable/, built on Anndata, is an one-stop package for single cell data analysis, including preprocessing, cell clustering, data visualization, etc. Scanpy utilizes the efficient methods and data structures aforementioned, and has incorporated other essential scRNA-seq data processing steps, e.g., normalization, quality control, batch effect removal, and clustering. With these features, Scanpy has become one of the most popular packages for scRNA-seq data analysis.

Though the use of efficient methods, algorithms, and proVRAMming, data analysis with Scanpy remains slow due to the extremely high dimensionality of single-cell RNA sequencing (scRNA-seq) data. Since scRNA-seq data is typically stored in matrix format, a natural approach is to leverage GPUs for parallel computing, as a single GPU card contains thousands of computational units, or CUDA cores. To address this, the scverse core team developed rapids-singlecell[7], a GPU-accelerated scRNA-seq analysis package that improves performance by more than 10 times compared to Scanpy. While rapids-singlecell was initially limited to datasets with fewer than 1 million cells, recent updates have expanded its capabilities to support massive-scale datasets using multiple GPUs via Dask, as well as out-of-core execution. This allows rapids-singlecell to efficiently handle extremely large datasets, beyond the constraints of GPU memory. For instance, PCA on 10 million cells can now be completed in under 10 seconds on a DGX system that designed by NVIDIA specifically for deep learning and artificial intelligence (AI) workloads.

Despite its improvements, rapids-singlecell still has limitations. While it supports multi-GPU configurations with Dask arrays for large datasets, it remains inaccessible to independent researchers who typically lack multiple GPUs or large-scale computing resources. Additionally, its out-of-core computing support, though promising, often faces issues with the RAPIDS Memory Manager (RMM), resulting in crashes or stalls. Furthermore, the Harmony algorithm, crucial for batch correction, suffers from high memory consumption and lacks multi-GPU support, limiting its scalability for datasets with many batches.

These limitations impact rapids-singlecell’s scalability, particularly for researchers with constrained resources or datasets that require memory-intensive algorithms like Harmony.

In this work, we introduce ScaleSC, a GPU-accelerated package developed on top of rapids-singlecell and Scanpy, designed to deliver their superior performance to resource-limited users. ScaleSC enables the analysis of large datasets on a single GPU, overcoming the memory constraints of rapids-singlecell. It delivers 20 to 100 times faster performance than Scanpy and can efficiently handle datasets with up to 20 to 40 million cells. Additionally, ScaleSC offers exceptional capabilities in marker gene identification and cluster merging.

A more detailed explanation of ScaleSC can be found in Section 2.

## Methods

Building on the foundation of rapids-singlecell, which already implemented most of the essential scRNA-seq data processing steps in the GPU version, we developed ScaleSC with several improvements, including 1) fixing the discrepancy of results between Scanpy and rapids-singlecell, 2) solving the scalability bottlenecks, and 3) optimizing algorithms.

### Discrepancy Removal

The output results are different between Scanpy and rapids-singlecell. Scanpy uses CPU-based scientific computing libraries like *NumPy, SciPy*, and *Sklearn*, while rapids-singlecell is mainly based on the GPU-accelerated *Rapids* AI ecosystem. Scanpy and rapids-singlecell implement the same algorithm slightly differently in order to optimize the different computational architectures between CPU and GPU. We refer to this difference as “system variance”. In addition to system variance, floating-point errors also contribute to differences in results, and we refer this as “numerical variance”. Although neither “system variance” nor “numerical variance” is incorrect, the difference in results leads to inconsistencies in data processing, posing challenges for reproducibility. Numerical variance is unavoidable but less significant compared to system variance and can be mitigated by increasing precision settings. This section focuses primarily on how ScaleSC eliminates system variance.

#### PCA discrepancy

Principal component analysis (PCA) is a critical step in single-cell RNA-seq pipeline, which efficiently projects high-dimensional single-cell count data into low-dimensional space for proceeding downstream analysis. PCA typically consists of two steps:

##### Standardization

The count value of each gene in each cell *X*_*ij*_ is normalized to a mean of 0 and a deviation of 1 as follows,

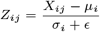

where *Z*_*ij*_ is the normalized count of gene *i* in cell *j, µ*_*i*_ is the mean count of gene *i* over all cells, *σ*_*i*_ is the standard deviation of gene *i* over all cells, *ϵ* is a constant to avoid divided by zero.

##### Eigen-Decomposition

An Eigen-Decomposition is performed on the covariance matrix. The eigenvectors represent the directions (principal components), and the eigenvalues represent the magnitude of the variance along those directions.

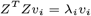

where *v*_*i*_ is the *i*-th principal component and *λ*_*i*_ is the corresponding eigenvalue of the *i*-th eigenvector.

The differences are raised by the sign of eigenvectors that is not uniquely defined, flipping the signs does not affect their mathematical validity, but this can result in a different visualization and make the following analysis not reproducible, for example, batch removal algorithm *Harmony* [8] takes principal components as inputs. As what we found, *Scanpy* internally calls PCA function from *Sklearn*, which aligns the direction by flipping the sign of *v*_*i*_ such that the largest absolute value in *v*_*i*_ is always positive, the correction can be formulated as below.

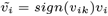

where *k* is the index of the largest absolute value. Instead of calling PCA from *Sklearn, rapids-singlecell* relies on *Rapids* suite that doesn’t correct it automatically, which makes the discrepancy appear. To further address this issue, one more step for correction of principal components is introduced in *ScaleSC*.

#### Harmony discrepancy

*Harmony* is a powerful method designed to integrate single-cell datasets from multiple batches or experiments while preserving the biological variation and removing batch effects. It operates on low-dimensional embeddings, typically generated by PCA, then initializes clusters by KMeans based on embeddings and repeatedly adjusts them until convergence.

We found the discrepancy arises from k-means clustering and randomness caused by Random Number Generator (RNG) even when the random seed remains unchanged. More specifically, CPU typically generates random numbers sequentially using algorithms like Linear Congruential Generators (LCG), and is deterministic when initialized with the same seed, while GPU generates random numbers in parallel without any guarantee of ordering. This property leads to the uncertainty of algorithms involving randomness, like KMeans. To ensure ScaleSC obtains the same initial clusters, the starting points of KMeans need to be matched up, we modified the original Python implementation *harmonypy* in two ways, The first approach is replacing GPU implementation of k-means with CPU implementation entirely, this minimizes the code changes and we call it 1-step modification. The second approach is switching the calculation of starting centroids to CPU then feed them to GPU-accelerated k-means, we found this way can maximally leverage GPU capacity compared to “1-step” modification, and this refers to 2-step modification.

#### Neighbor discrepancy

Finding the nearest neighbors based on the *Harmony* outputs is essential for subsequent Leiden clustering. We aim to construct clusters using the same neighborhood graph and cell connectivities. However, we observe that Scanpy and rapids-singlecell generate two distinct graphs from the same input. Upon a detailed review of both Scanpy and rapids-singlecell implementations, we identified differences in their nearest neighbor algorithms. rapids-singlecell utilizes an exact k-Nearest Neighbor (k-NN) algorithm, implemented in parallel via *cuML*, while Scanpy employs an approximate k-NN method using NNdescent[9], an algorithm designed for approximating nearest neighbor searches in high-dimensional spaces. To validate this distinction, we forced Scanpy to use the exact k-NN algorithm and confirmed that the results aligned with those produced by rapids-singlecell. Consequently, ScaleSC will call rapids-singlecell internally, using exact k-NN rather than the approximation.

By incorporating the improvements described above, ScaleSC can replicate Scanpy’s results in most steps. However, even with these discrepancies resolved, certain steps involving stochastic algorithms, such as Leiden, UMAP, and t-SNE, may not consistently yield identical results. These variations, primarily introduced during visualization, do not impact the conclusions of quantitative analyses.

### Scalability

Frequent interaction with the count matrix can’t be avoided in the single-cell RNA-seq preprocessing pipeline, loading the whole matrix into memory during runtime is infeasible. Due to its high sparsity (typically *>* 90% non-zero values), equivalent sparse formats are preferred for storage and computing on both CPU and GPU. However, there are two bottlenecks prevent us from actually using GPU:

#### VRAM Bottleneck

Most of high-end GPUs in the market have limited memory. Even like the most advanced NVIDIA A100 has only 80*G*, which is insufficient for one-time loading for dataset even with sparse matrix supported.

#### Int32 Bottleneck

*CuPy* as the backend of sparse matrix on GPU, one notable limitation is the integer type used for indexing, which is commonly 32-bit integers, indicating it can only store non-zero elements up to 2^31^ − 1, or approximately 2.1 billion. This is not enough for extremely large single-cell dataset. For example, a dataset with 5 *×* 10^6^ cells, 2 *×* 10^4^ genes, and a sparsity of 0.9, the number of non-zero elements is much larger than that upper limit. Currently, there is no solution to solve it directly.

To work around with extremely large dataset, we adopt the chunking approach to split the dataset into chunks along cells, so that each data chunk won’t reach the Int32 bottleneck. In addition, this also gives us flexibility to process data chunks separately across devices, which indicates there is a potential way to process arbitrarily large data.

#### ScaleSC overview

Next, we build ScaleSC on top of Scanpy and rapids-singlecell so that our method can seamlessly scale to tens of millions of cells with GPU acceleration. Like Scanpy and rapids-singlecell, ScaleSC implements a series of necessary methods in the classical single-cell RNA-seq pipeline, including steps shown in Figure. 1:

**Fig. 1.**
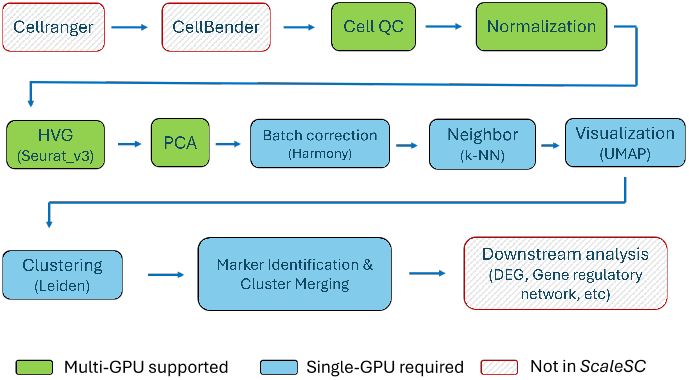
The Workflow of standard scRNA-seq data processing.

1. Cell QC: Filter out low-quality cells and genes according to the number of counts.
2. Normalization: Perform log1p normalization on counts by log(1 + *X*), where *X* is the count value.
3. HVG: Choose highly variable genes using Seurat v3 [5].
4. PCA: Dimensionality reduction using PCA.
5. Batch Correction: Invoke *Harmony* to remove batch effects.
6. Neighbor: Use *k*-NN to find the *k* nearest neighbors to build the neighborhood graph.
7. Visualization: Call UMAP on PCA-reduced matrix for better visualization.
8. Clustering: Call Leiden clustering.
9. Marker Identification: Identify cluster-specific genes.
10. Cluster Merging: Clusters are merged to eliminate those that are deemed non-informative, based on the markers detected during the analysis.

Since ScaleSC conducts data loading and preprocessing in data chunks, we developed a chunked-data class dedicated to chunked data loading and computation. This reader supports three different modes for loading data under various scenarios, depending on the user’s GPU and CPU size. The three data loading modes are described in the next subsections.

**Fig. 2.**
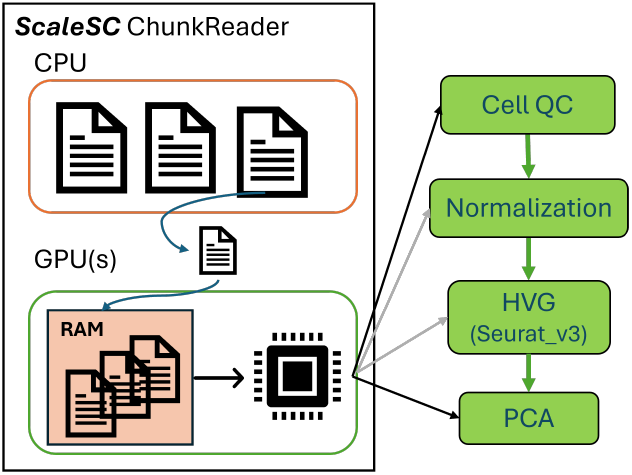
The flowchart Chunk-based data loading and preprocessing.

Time-efficient but high memory consumption In the implemented architecture, ScaleSC employs an optimized memory management strategy by pre-loading all data chunks into GPU memory (VRAM). The processing pipeline operates iteratively, where individual chunks are sequentially processed, followed by automatic result aggregation. This approach eliminates the computational overhead associated with CPU-GPU data transfer during processing, as all data resides in VRAM, thereby maximizing computational efficiency. However, this implementation is constrained by available VRAM capacity, limiting the maximum processable dataset size. To overcome this limitation and accommodate larger datasets, we introduced multi-GPU support for distributed chunk storage. This extension leverages the high-bandwidth of GPU-to-GPU data transfer capabilities through NVSwitch, which significantly outperforms CPU-to-GPU transfer rates. Notably, while data chunks are distributed across multiple GPUs for storage, computation remains centralized on a single GPU to maintain processing efficiency and algorithmic consistency.

#### Balanced on both time and memory

In contrast to the GPU-centric approach, ScaleSC implements an alternative mode where data chunks are pre-loaded and maintained in CPU memory rather than GPU VRAM. In this implementation, the processing pipeline transfers individual chunks sequentially to GPU for computation, followed by result retrieval back to the CPU memory. While this approach introduces additional overhead from CPU-GPU communication, it circumvents GPU memory constraints by leveraging typically larger CPU memory capacity. This implementation’s performance is primarily bounded by available CPU memory and includes latency from device communication overhead. However, empirical evidence suggests this mode provides robust performance across a wide range of real-world applications, offering a practical balance between computational efficiency and memory management.

#### Memory efficient but time-consuming

For resource-constrained environments, ScaleSC implements a disk-based processing mode that minimizes memory requirements. In this implementation, data chunks are read directly from disk storage on demand, rather than being pre-loaded into either CPU or GPU memory. While this approach introduces significant input/output overhead between disk and host memory, it provides minimal memory footprint at the cost of increased processing time. This mode’s primary advantage lies in its theoretical capacity to process datasets of arbitrary size, constrained only by available disk space rather than system memory. The trade-off between processing speed and memory utilization makes this implementation particularly valuable for environments with limited computational resources, ensuring ScaleSC’s accessibility across diverse computational infrastructures.

### Optimizations in ScaleSC

This section examines the architectural modifications implemented in ScaleSC to efficiently process chunked data structures. We detail the algorithmic adaptations necessary for chunked based computation and subsequently explore enhancements to the Harmony batch correction algorithm to accommodate substantially larger batch sizes. These modifications are essential for maintaining computational efficiency while processing large-scale single-cell datasets.

#### Cell QC

Within each individual chunk, a subset of cells is retained, while all genes are preserved. This approach facilitates the filtering of cells based on the number of gene counts. In contrast, selecting genes based on the number of cells presents a challenge, as other chunks are not accessible when processing a specific chunk. To address this, an array is maintained to track the number of cells associated with each gene, which is updated incrementally as new chunks are processed. Ultimately, a finalized list of cells and genes is generated following a single iteration over all chunks. This procedure requires traversing all chunks once to ensure the completeness of the filtering process.

#### HVG

This section outlines the implementation of the HVG selection step, following the methodology described in the Seurat v3 paper [5]. The workflow can be summarized as follows: *N* represents the number of cells and *M* represents the number of genes:

1. The means and variances for each gene are computed using the raw count matrix. This process requires a single iteration over all data chunks to calculate the means, denoted as *µ*_*i*_, for the *i*th gene, and the sample variances, denoted as *σ*_*i*_, for the *i*th gene (using *N*−1 as the divisor). It is important to note that using population variance instead of sample variance will introduce discrepancies.

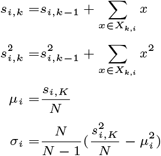

where *K* is number of chunks, *X*_*k,i*_ is the *i*th gene column in the *k*-th chunk, *x* is the count value, *s*_*i,k*_ is sum of count values in the first *k* chunks, 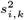 is the sum of squared count values in the first *k* chunks.
2. Loess Regression in Logarithmic Space: The locally weighted regression (LOESS) is performed by using the logarithm of means as the predictor and the logarithm of variances as the response variable. While a GPU implementation is not currently available, the method is sufficiently fast and efficient when executed on the CPU.
3. Normalization and Clipping: Normalize the raw counts using the fitted standard deviation and means for each gene, and clip any values exceeding 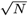.

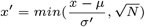

where *x*^*′*^ is the normalized value, *µ* is the mean of the gene *i* before normalization, *σ* is the fitted standard deviation in Step(2).
4. Repeat Step(1) to recalculate the variance of normalized data, then mark genes with the K-largest variances as HVGs.

In HVG, two loops of all chunks are needed: the first loop obtains mean and variance of raw data; the second loop normalizes data with regressed variances.

#### PCA

Given the chunked nature of the data, obtaining eigenvectors through Singular Value Decomposition (SVD) is not straightforward. However, it is considerably easier to derive them through the use of a zero-centered covariance matrix.

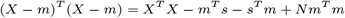

where *X* ∈ *R*^*N×M*^ represents the entire matrix, *m* ∈ *R*^*M*^ is a vector of means, *s* ∈ *R*^*M*^ is a vector representing the sum of count values. We can calculate *X*^*T*^ *X* by simply adding up all chunks.

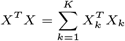

where *X*_*k*_ is the *k*-th chunk, and 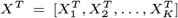. Based on the calculated zero-centered covariance matrix, the principal components can be easily obtained through eigen-decomposition. Similar to the High Variance Genes (HVG) step, ScaleSC processes all chunks twice.

#### Batch Correction

*Harmony* [8] is a widely used algorithm for batch effects removal. However, during these iterative calculations, the algorithm generates and stores numerous intermediate variables, leading to substantial memory consumption. For datasets with approximately 13 million cells and over 1,000 samples, the original Harmony implementation encounters significant memory issues, often resulting in out-of-memory (OOM) errors. The memory requirement for such datasets exceeds 220 GB, highlighting a critical limitation in its scalability for large-scale single-cell analyses.

From Table 4 in supplementary, the obvious bottleneck is *B*. We can observe that when *B* increases, it becomes impossible to hold even such a single large matrix under 32-bit float. However, Φ, Φ_*moe*_ and *Phi*_*Rk*_ are extremely sparse containing about *N* non-zero values. If we convert them to sparse matrix, a significant amount of memory can be saved. Based on this strategy, a (*B, N*) dense matrix can be converted to a sparse matrix in only *O*(*N*) space (∼ 0.2*G*). All algebraic operations involving converting matrices need to be adjusted accordingly as well.

The original script also used dummy variables to create the matrix Φ, which takes a lot of memory on CPU as well. To further optimize memory performance, we generate a sparse dummy matrix directly instead of a huge dense matrix. Besides those sparse matrices, the original script keeps some unused matrices in memory. We removed them and further optimized the script to save more memory.

After conversion and cleaning, it still fails on 13*M* data. The reason is *Harmony* produces an extremely large intermediate values when it estimates MoE model parameters as following:

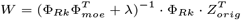

A detailed explanation of Φ_*Rk*_, Φ_*moe*_, and *Z*_*orig*_ is provided in the Table 4 in the supplementary material. Multiplication of 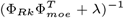 and Φ_*Rk*_ results in an intermediate matrix with shape as (*B, N*), which is approximately 64*G* memory .space allocated by GPU for temporary storage. A simple trick can be observed to avoid such large temporary storage appeared during calculations: 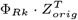 won’t take too much space so that can be computed first, then multiply by 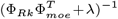.

### Marker Gene Identification

Cell type annotation plays a crucial role in cell type decomposition analysis and understanding biological processes, such as pathways enriched in certain cell types. However, accurately annotating cell types remains a challenge. There are two main approaches: de novo annotation, which relies on marker genes, and reference mapping, which transfers annotations from well-annotated datasets. Marker genes are those that are highly expressed in a specific cluster while showing low expression in others. In Scanpy, marker genes are identified using the function “Scanpy.tl.rank genes groups”, which employs methods like t-test. However, the identified markers often lack accuracy and specificity (see Figure 3).

**Fig. 3.**
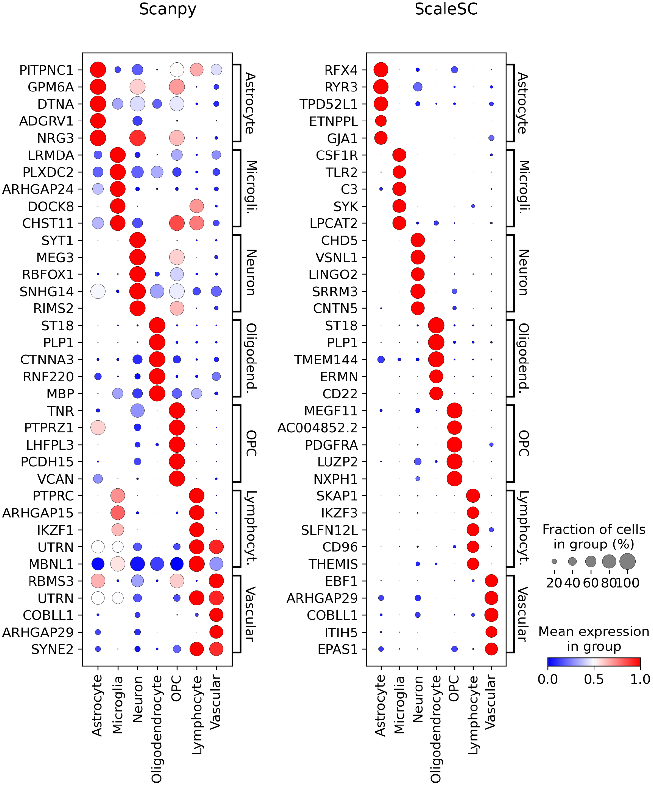
Expression of gene markers between Scanpy and ScaleSC is compared using a dataset with 0.6M cells. The left panel displays the markers detected by Scanpy, while the right panel shows the markers identified by ScaleSC. Rows correspond to the markers, and columns represent the clusters. Brackets indicate cluster-specific markers within each cluster, with larger circles denoting a higher proportion of the marker within the cluster. The expression values are standardized to a range between 0 and 1.

NSForest [10] offers a more advanced approach by integrating Random Forest algorithms with network structure analysis. Despite its advantages, its computational complexity has limited its application to smaller datasets, making it impractical for large-scale single-cell analyses. To overcome this, we resolved key memory and computational bottlenecks and developed a GPU-accelerated version. By leveraging modern GPUs’ parallel computing capabilities, our optimized approach enables the analysis of datasets with millions of cells while preserving NSForest’s strong feature selection performance. A comparison of results between Scanpy and ScaleSC is shown in Figure 3.

## Results

### Time efficiency

To comprehensively evaluate the performance of Scanpy, rapids-singlecell, and ScaleSC, we selected datasets with three different scales: small, medium, and large. Each method has been tested across all datasets, from data loading to Leiden clustering, with the time taken for each step recorded for comparison. All tests were conducted on a system running Rocky Linux 8.10, equipped with 1 TB of CPU RAM and a single NVIDIA A100 GPU.

For the small dataset, we utilized the 70K human lung dataset[11], which contains 65,662 cells and 26,485 genes. All three methods were able to run on this dataset. As shown in the Table 1, both ScaleSC and rapids-singlecell exhibit exceptional performance, achieving a 15.4x speedup.

**Table 1.** The time cost comparison between Scanpy, rapids-singlecell, and ScaleSC on the 70K human lung data [11].

| Step | Scanpy(sec) | rapids-singlecell(sec) | ScaleSC(sec) | Speedup |
| --- | --- | --- | --- | --- |
| Data loading | 0.9 | 0.9 | 1.4 | 0.7 |
| QC | 7.5 | 0.1 | 0.1 | 75.0 |
| Normalization | 1.1 | 0.0 | 0.1 | 11 |
| HVG | 10.5 | 0.3 | 0.2 | 52.5 |
| PCA | 14.8 | 0.3 | 0.8 | 18.5 |
| Harmony | 131.4 | 7.8 | 22.0 | 6.0 |
| Neighbors | 153.5 | 0.5 | 0.7 | 218.6 |
| UMAP | 87.5 | 0.2 | 0.2 | 437.0 |
| Leiden clustering | 16.2 | 2.7 | 2.0 | 8.1 |
| **All** | **423.4** | **12.8** | **27.5** | **15.4** |

For the medium-scale dataset, we used the 1.3M mouse brain dataset[4]. rapids-singlecell is not feasible for this dataset due to exceeding the CuPy’s int32 limitation. ScaleSC completed the task in 2 minutes, while Scanpy requires 4.5 hours to process the entire dataset as shown in Table 2.

**Table 2.** The time cost comparison between Scanpy and ScaleSC on the 1.3M mouse brain data [4].

| Step | Scanpy(sec) | ScaleSC(sec) | Speedup |
| --- | --- | --- | --- |
| Data loading | 26.0 | 16.1 | 1.6 |
| QC | 271.6 | 4.9 | 55.6 |
| Normalization | 21.3 | 2.0 | 10.7 |
| HVG | 312.5 | 8.5 | 37.0 |
| PCA | 269.6 | 21.7 | 12.4 |
| Harmony | 8818.1 | 23.5 | 375.5 |
| Neighbors | 336.5 | 34.1 | 9.9 |
| UMAP | 2404.3 | 4.4 | 546.8 |
| Leiden clustering | 3793.2 | 10.9 | 346.6 |
| **All** | **16253.1** | **126.0** | **128.9** |

Note: rapids-singlecell is not compared here due to CuPy’s int32 limitation.

To assess the performance of ScaleSC on extremely large datasets, we generated a simulated dataset containing 13 million cells based on the original 1.3 million-cell mouse brain dataset. This expanded dataset was created by generating simulated samples from the original samples, with each sample being simulated nine times. For a given sample, new cells were generated for each cluster by modeling the distribution of gene expression within that cluster. The results, presented in Table 3, demonstrate that, with the optimizations implemented, ScaleSC completed the analysis in under one hour while utilizing approximately 60 GB of GPU memory. This performance indicates that ScaleSC is highly efficient for handling large-scale datasets and holds promise for scaling up to 20 million cells on a single A100 GPU. In contrast, both Scanpy and rapids-singlecell failed to process such large datasets.

**Table 3.** The time cost of ScaleSC on a simulated data with over 13M cells.

| Step | ScaleSC(sec) |
| --- | --- |
| Data loading | 405 |
| QC | 686 |
| Normalization | 56 |
| HVG | 419 |
| PCA | 622 |
| Harmony | 628 |
| Neighbors | 163 |
| UMAP | 27 |
| Leiden clustering | 49 |
| **All** | **3055** |

Note: Both Scanpy and rapids-singlecell are infeasible on this dataset.

### Gene Marker Identification

We evaluated the performance of ScaleSC against Scanpy using a medium-scale single-cell RNA sequencing dataset comprising 661, 789 cells and 36, 341 genes. To assess baseline performance characteristics, the comparison was conducted without preliminary feature selection through highly variable gene filtering. rapids-singlecell was excluded from this comparison due to its methodological similarity to Scanpy’s implementation.

As shown in Fig. 3, the marker genes identified by ScaleSC demonstrated superior biological relevance, characterized by both increased cell-type specificity and elevated expression levels. Specifically, these markers exhibited higher group-wise cell fraction representation and enhanced expression intensity compared to traditional approaches, suggesting improved capacity for cell-type discrimination. Additional comparisons on another two datasets are shown in supplementary.

## Conclusion

The growing scale of single-cell RNA-seq data presents significant challenges in terms of processing speed and memory capacity. In this work, we introduced ScaleSC, a GPU-accelerated solution designed specifically to address these challenges for users with limited computing resources, such as those with a single GPU or without deep familiarity with GPU ecosystems. By building upon the powerful foundations of CuPy, Scanpy, and rapids-singlecell, ScaleSC delivers over a 20x speedup in processing, overcoming memory bottlenecks and enabling the analysis of datasets with 10–40 million cells and over 1,000 batches on a single A100 GPU. In addition to improving scalability, ScaleSC ensures consistent results by resolving discrepancies between GPU and CPU algorithm implementations. The package also includes GPU-optimized tools for advanced tasks like marker gene identification and cluster merging, providing more efficient solutions for better marker identification. Designed for ease of use, ScaleSC integrates seamlessly into existing Scanpy and rapids-singlecell workflows, requiring minimal adaptation from users. With its substantial improvements in speed, scalability, and accessibility, ScaleSC offers significant benefits to the single-cell RNA-seq community, particularly for those with limited computational resources, providing an accessible and high-performance solution for large-scale data analysis.

## Author contributions statement

- W.H. and H.Z. Conceptualization, Methodology, Data Analysis.
- H.Z. Software Development.
- W.H., H.Z., Y.H.S., S.C., J.G., Z.O., Y.M. and B.Z. Writing & Review.

All authors reviewed and approved the final manuscript.

## Acknowledgments

The authors thank the anonymous reviewers for their valuable suggestions.

## Supplementary Material

**Table 4.** Some large variables produced during calculation (13 million cells), *d* denotes the dimension of PC loadings, *B* denotes the number of samples, *C* denotes the number of clusters (up to 100).

| Variable | Description | Shape | Size |
| --- | --- | --- | --- |
| Z_orig | PCA input | $(d, N)$ | $2.5G$ |
| Z_cos | L2 normalization of input, cosine distances | $(d, N)$ | $2.5G$ |
| Z_corr | Corrected embeddings | $(d, N)$ | $2.5G$ |
| Phi | The one-hot encoded batch assignment matrix. | $(B, N)$ | $64G$ |
| Phi_moe | The one-hot batch assignment with bias in MoE estimation | $(B + 1, N)$ | $64G$ |
| Phi_Rk | The one-hot batch assignment for a given cluster | $(B + 1, N)$ | $64G$ |
| R | the cluster assignment matrix | $(C, N)$ | $5G$ |
| dist_mat | Distance matrix | $(C, N)$ | $5G$ |
| scale_dist | Scaled distance matrix | $(C, N)$ | $5G$ |

**Fig. 4.**
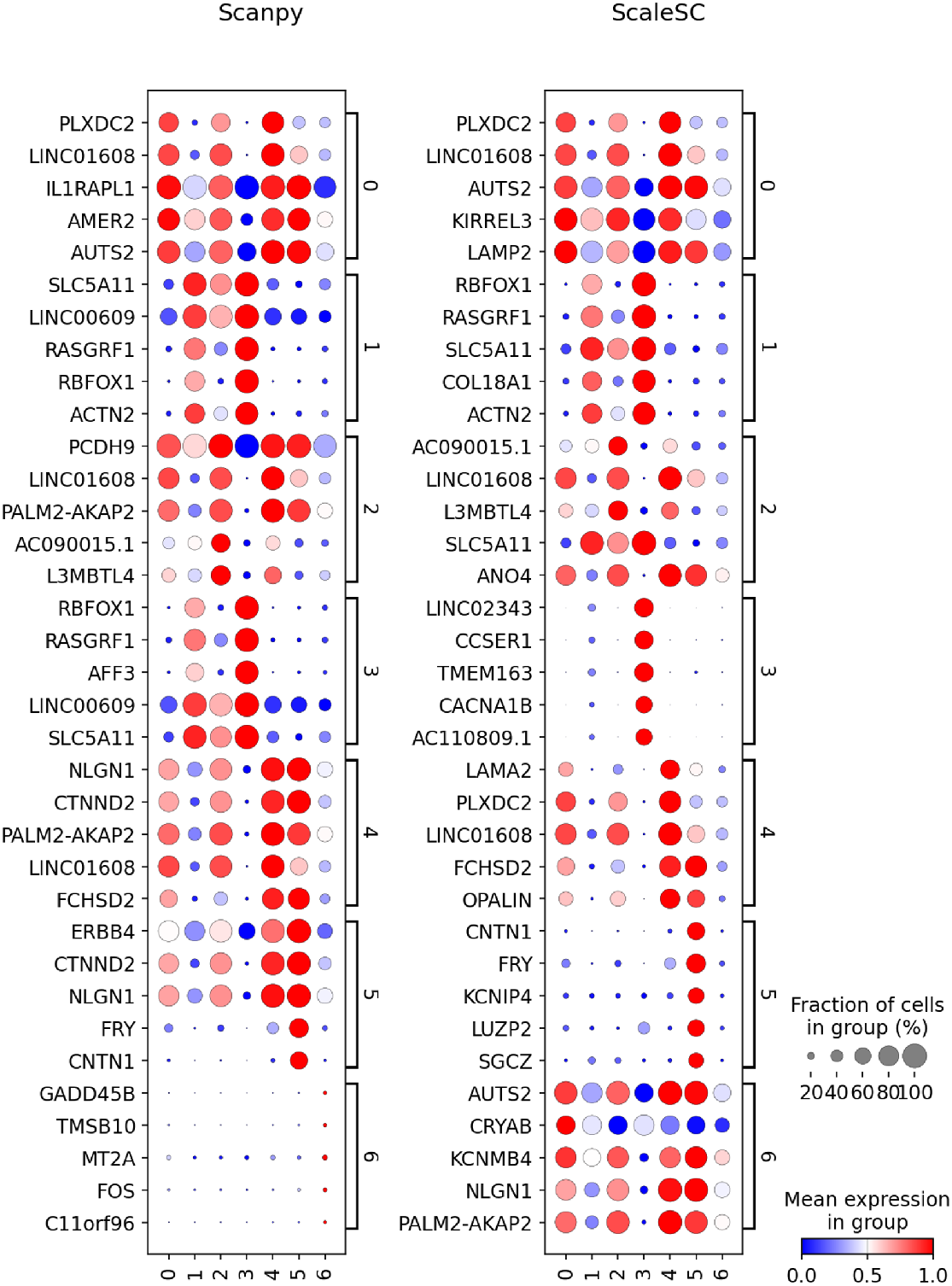
Expression of gene markers between Scanpy and ScaleSC is compared using a dataset with 18, 262 cells and 36, 341 genes. The left panel displays the markers detected by Scanpy, while the right panel shows the markers identified by ScaleSC. Rows correspond to the markers, and columns represent the clusters. Brackets indicate cluster-specific markers within each cluster, with larger circles denoting a higher proportion of the marker within the cluster. The expression values are standardized to a range between 0 and 1.

**Fig. 5.**
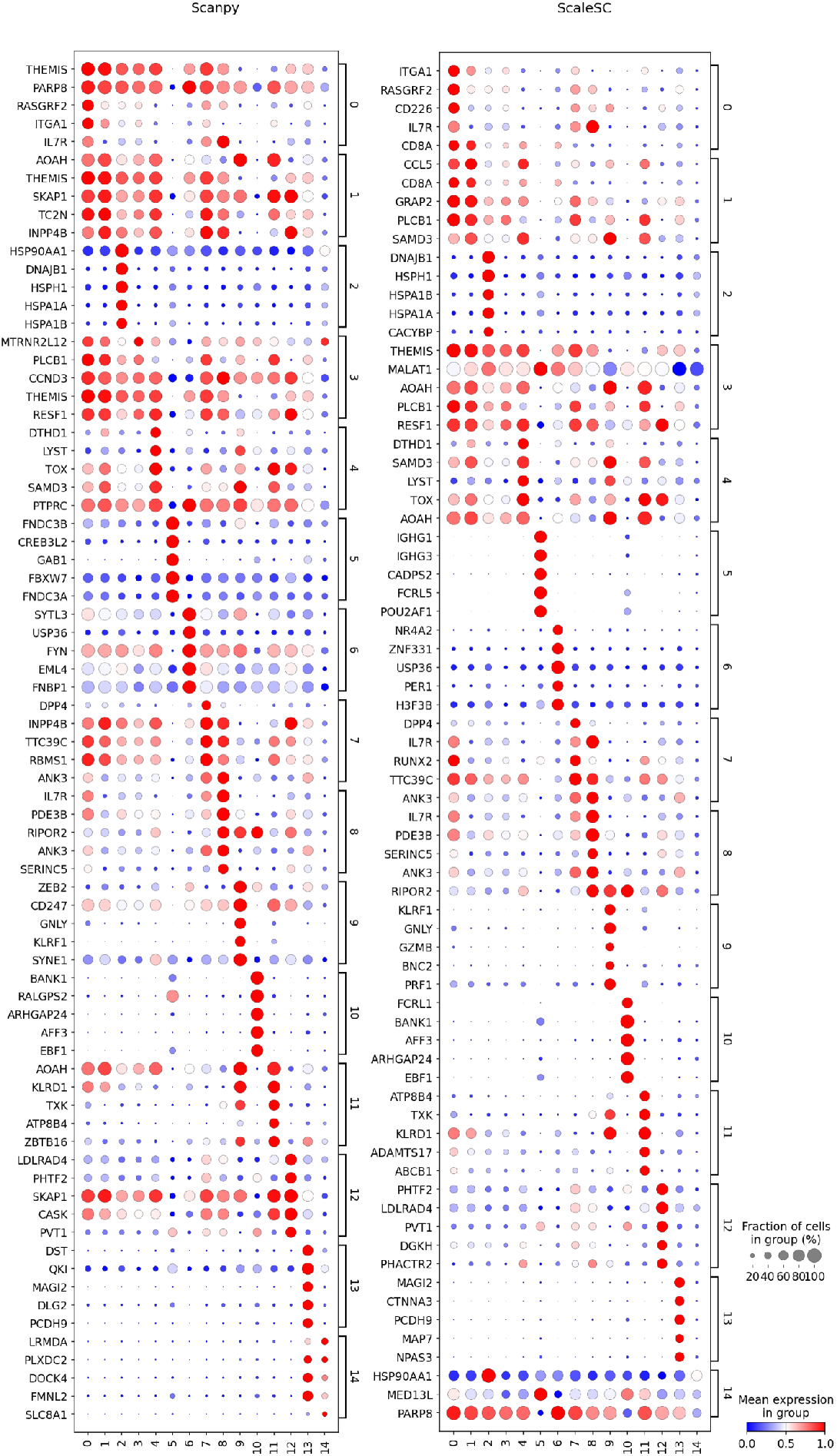
Expression of gene markers between Scanpy and ScaleSC is compared using a dataset with 0.5M cells and 36, 341 genes. The left panel displays the markers detected by Scanpy, while the right panel shows the markers identified by ScaleSC. Rows correspond to the markers, and columns represent the clusters. Brackets indicate cluster-specific markers within each cluster, with larger circles denoting a higher proportion of the marker within the cluster. The expression values are standardized to a range between 0 and 1.

